# Frataxin-deficient human brain microvascular endothelial cells lose polymerized actin and are paracellularly permeable –implications for blood-brain barrier integrity in Friedreich’s Ataxia

**DOI:** 10.1101/2023.02.09.527936

**Authors:** Frances M. Smith, Daniel J Kosman

**Affiliations:** Department of Biochemistry, Jacobs School of Medicine and Biomedical Sciences, The University of New York at Buffalo

**Keywords:** Friedreich’s Ataxia, blood-brain barrier, barrier permeability, filamentous actin, neuropathology.

## Abstract

**Background:** Friedreich’s Ataxia (FRDA) is the most prevalent inherited ataxia; the disease results from loss of Frataxin, an essential mitochondrial iron trafficking protein. FRDA presents as neurodegeneration of the dorsal root ganglion and cerebellar dentate nuclei, followed by brain iron accumulation in the latter. End stage disease includes cardiac fibrosis that contributes to hypertrophic cardiomyopathy. The microvasculature plays an essential barrier role in both the brain and heart, thus an investigation of this tissue system in FRDA is essential to the delineation of the cellular dysfunction in this genetic disorder. Here, we investigate brain microvascular endothelial cell integrity in FRDA in a model of the blood-brain barrier (BBB).

**Methods:** We used lentiviral mediated shRNA delivery to generate a novel FRDA model in immortalized human brain microvascular endothelial cells (hBMVEC) that compose the microcapillaries of the BBB. We verified known cellular pathophysiologies of FXN knockdown including increased oxidative stress, loss of energy metabolism, and increased cell size. Furthermore, we investigated cytoskeletal architecture including the abundance and organization of filamentous actin, and barrier physiology *via* transendothelial electrical resistance and fluorescent tracer flux.

**Results:** shFXN hBMVEC display the known FRDA cell morbidity including increased oxidative stress, decreased energy metabolism, and an increase in cell size. We demonstrate that shFXN hBMVEC have less overall filamentous actin, and that filamentous actin is lost at the cell membrane and cortical actin ring. Consistent with loss of cytoskeletal structure and anchorage, we found decreased barrier strength and increased paracellular tracer flux in the shFXN hBMVEC transwell model.

**Conclusion:** We identified that insufficient FXN levels in the hBMVEC BBB model causes changes in cytoskeletal architecture and increased barrier permeability, cell pathologies that may be related to patient brain iron accumulation, neuroinflammation, neurodegeneration, and stroke. Our findings implicate other barrier cells, *e.g.,* the cardiac microvasculature, likely contributory also to disease pathology in FRDA.

## Introduction

Friedreich’s Ataxia (FRDA) is the most prevalent inherited ataxia, affecting ∼1:50,000 US citizens ^1^. FRDA is diagnosed around the age of 15, with∼20 years of degenerating quality of life before patient death around the age of 37 often due to cardiac fibrosis contributory to hypertrophic cardiomyopathy ^2, 3^. FRDA is caused by GAA expansion repeats within the first intron of the Frataxin (*FXN*) gene, leading to replication stress and transcriptional repression which thereby decreases protein production ^4, 5^. FXN is an essential protein localized to the mitochondrial matrix space, assisting in iron incorporation into iron-sulfur clusters (ISC) and heme centers ^6^. In mice, the homozygous *fxn*/*fxn^-^* genotype is embryonic lethal. ISC and iron porphyrin prosthetic groups support diverse enzymatic functions including DNA repair, iron homeostatic dynamics, and importantly, energy metabolism. Thus, FRDA is a metabolic disease, as patients experience severe mitochondrial defects, decreased electron transport chain (ETC) complex expression, depressed oxidative energy metabolism, diabetes mellitus type-II, and buildup of blood lactate ^7–12^.

Patient neurologic symptoms include atrophy of the cerebellar dentate nuclei (CDN) and dorsal root ganglion (DRG); progressive iron accumulation occurs in the former ^13, 14^. Degeneration of these high-velocity cognitive processing centers confers loss of sensory-motor and limb control, and ataxic gait. The CDN is suggested to be susceptible to iron accumulation due to its naturally-enriched iron content, but FRDA brain-iron pathology remains poorly understood ^13^. Recent longitudinal studies indicate that CDN pathology in FRDA starts with a burst of neurodegeneration, followed by progressive brain iron accumulation ^14^. That the brain iron accumulation occurs longitudinally with disease progression indicates that it is an active deposition process, supported by iron flux through (*transcellularly*) or around (*paracellularly)* brain microvascular endothelial cells.

The loss of FXN iron chaperoning capacity in FRDA cells results in intra-mitochondrial iron accumulation, a process that generates reactive oxygen species (ROS). This is further compounded by loss of antioxidant signaling via decreased activity of Nrf2, a major antioxidant transcription factor ^12, 15, 16^. In addition, levels of free glutathione, a redox regulator, is found to be significantly depleted in FRDA patient-isolated fibroblasts. This is co-incident with glutathionylation of actin monomers, a process now known to be inhibitory of actin filament formation in both the rate and extent of polymerization ^17, 18^. This is an indirect modulation of the cytoskeleton in FRDA. There is direct modulation as well, including the transcriptional repression of PIP5K-1β due to the large GAA repeat expansion at the *FXN* locus; ***PIP5K1β*** flanks *FXN* on chromosome 9. PIP5K-1β is functionally linked to actin cytoskeletal dynamics, and its loss in FRDA patient fibroblasts leads to altered cell spreading and actin remodeling dynamics ^19^. We hypothesized that a similar degradation of the actin network in FRDA would compromise the tight junction network between hBMVEC essential to the barrier function of these cells.

The blood-brain barrier (BBB) is a network of microvascular endothelial cells which form an impermeable barrier due to apically-most facing tight junctions (TJs), maintaining separation of the circulation and the brain interstitium. The BBB is selectively permeable *via* uptake and efflux transporters and vesicular trafficking, but paracellularly impermeable to molecules exceeding 400 Daltons ^20^. Actin filaments anchor the transmembrane TJ proteins (claudins and occludins) *via* the cytosolic protein zona occludens (ZO) family members ^21, 22^. The transmembrane TJ proteins then form homodimers with their same family members on adjacent cells to maintain paracellular impermeability ^23^. Based on the known actin pathology in FRDA patient cells, we hypothesized that the barrier properties of the endothelial cells of the BBB are negatively affected by cytoskeletal alterations associated with FXN loss ^17, 24^. While FXN expression is high in brain homogenate, the presence or role in brain microvascular endothelial cells has not been investigated. We sought to extend FRDA research in this highly specialized and essential microvascular system.

BBB research is limited in the FRDA literature, and examination of barrier function could provide therapeutic targets for FRDA neurophysiology. Here we examine the vascular integrity in FXN-knockdown human brain microvascular endothelial cells (hBMVEC Following shRNA-mediated FXN knockdown, hBMVEC display a loss of filamentous actin at the cell membrane and paracellular permeability to a fluorescent tracer. Our findings suggest that investigation of vascular integrity in FRDA patients likely would open new therapeutic windows addressing brain solute influx and neuropathology in this disease.

## Methods

### Reagents

All chemical reagents are obtained from Sigma Aldrich unless otherwise stated. Lyophilized compounds are solubilized per manufacturer instructions. All protocols are performed per manufacturer instructions unless otherwise stated.

### Antibodies

Rabbit α-FXN (ThermoFisher PA5-13411), Goat α-rabbit:HRP (Cell Signaling Technologies 7074), Mouse α-TATA Binding Protein (ThermoFisher 49-1036), Goat α-mouse:HRP (Novus Biologicals NBP2-31347H), and Rabbit *α*-β-actin (Cell Signaling Technologies 8457).

### qPCR Primers

FXN-F (TGGAATGTCAAAAAGCAGAGTG), FXN-R (CCACTCCCAAAGGAGACATC), β-2-Microglobulin-F (GCTCGCGCTACTCTCTCTTT), β-2-Microglobulin-R (CGGATGGATGAAACCCAGACA)^25^, and β-actin-F (GGGTCAGAAGGATTCCTATG), β-actin-R (GGTCTCAAACATGATCTGGG).

### Cell culture

Wild-type human brain microvascular endothelial cells (hBMVEC) are an immortalized cell line, a gracious gift from Dr. Supriya Mahajan (University at Buffalo). Verification of proper cell behavior is provided ^26, 27^. hBMVEC were cultured in RPMI, 10% FBS (Gibco) and 10% Nuserum (Corning), passaged at confluency and used between passage numbers 7 and 15 in experimentation. shRNA-integrated cells are continued in culture under selective pressure at 7.5ug/ml puromycin in RPMI until plated for an experiment, at which all cells are in normal growth media. Cells were housed in a 37°C incubator, 5% CO_2_ injection, and constant humidity.

### Knockdown

shRNA was generated against FXN using the commercial pGIPZ system along with their empty vector (EV) backbone control (Horizon Discovery). Briefly, glycerol stocks were plated and selected on Ampicilin-containing LB agar at 30°C overnight. Single colonies were selected and grown in liquid culture for MaxiPrep (Omega EZNA) at 37°, shaking overnight. Plasmid DNA for shFXN, EV, and psPAX2 and pMD2.G packaging systems (gracious gifts from Didier Trono: Addgene plasmids 12260 and 12259 respectively) were eluted into TE elution buffer, and concentration and purity were quantified via NanodropOne (Invitrogen). HEK293T cells were plated at 5 million cells/10cm dish in DMEM +10% FBS and let attach overnight. The following morning, 15µg psPAX2, 6µg pMD2.G, and 20µg of each shRNA or EV plasmid DNA were combined with CaCl2 and HBS for Calcium Phosphate transfection as described ^28^. Briefly, the DNA mixture was incubated at room temperature for 15 minutes then added dropwise to the HEK293T cells. Media was changed 6 hours following transfection to normal growth media. Virus-containing media was collected at 48- and 72-hours following transfection, filtered through 0.22µM PES filter, and concentrated using a sucrose-gradient ultracentrifugation.

The final viral pellet was solubilized in RPMI +10% FBS, 10%Nuserum. 20µL of each virus (shFXN and EV) + 5µg/ml Polybrene was added to wild-type hBMVEC for 6 hours, then replaced with growth media. 2 days following transfection, shRNA-containing cells were selected for in growth media + 0.75µg/ml Puromycin. Cell populations were allowed to grow back to confluency with minimal media changes. Proper shRNA expression was verified by the presence of GFP expression using the ZOE fluorescent imager.

### Reverse transcription quantitative polymerase chain reaction (RT-qPCR)

hBMVEC were lysed in Trizol (Ambion Life Technologies) and RNA was isolated using Direct-Zol MiniPrep spin columns (Zymogen) per manufacturer instructions with additional RNAse-OUT (Invitrogen) treatment during DNA digestion. RNA eluted in RNAse-free water was quantified for both concentration and purity using the NanodropOne (Invitrogen). 400ng of RNA was reverse transcribed using qScript (QuantaBio), of which 15ng each was loaded into a qPCR plate with a master mix of 5µL iTaq SYBR Green (Biorad) and 300nM each forward and reverse primers for; FXN, β2M, and *β*-actin. The absence of gDNA was confirmed by cycling equimolar concentrations of RNA that had not been reverse transcribed. All qPCR reactions were cycled with the following conditions using the CFX96 Touch qPCR (Biorad); 95°C 30s, 95°C 5s, 51°C 60s, x40 cycles, 65-95°C 0.5s increment. Differential abundance of transcript was assessed using the - DDCt method of quantitation with *β*2M as the housekeeping gene, normalized to the values of the EVEC control.

### Western blotting

hBMVEC were lysed in RIPA buffer containing 4x protease inhibitor and kept on ice. Lysates were spun at 4°C for 15 minutes at 13,000RPM. Supernatant was separated from cell debris and protein quantified using the BCA method at 562 nm absorbance (Thermo Scientific). 20-25μg of protein was added to 1x Laemli-buffer containing 150mM dithiothreitol (DTT) for denaturation at 37°C for 30 minutes. Lysates were electrophoresed on a 12% bis-tris Bolt gel (ThermoFisher) alongside 2.5µL Magic Mark protein molecular weight marker (Thermo Fisher) and 3.5μl Dual Color Xtra standard protein ladder (BioRad) at 140V. The gels were then transferred to a water-activated nitrocellulose membrane at 1.3A to 25V for 7 minutes (BioRad TurboBlotter, Mixed Molecular Weight setting). Membranes were blocked in Every Blot Blocking Buffer (Biorad) for 10 minutes, rocking at room temperature. All primary antibodies were diluted in 1% milk and TBST, and rocked over the solution at 4°C overnight (*α*-FXN; 1:1000, *α*-TATA-binding protein; 3.2μg/ml, *α*-β-actin 1:5,000). The next day, the membrane was washed thrice in 10-minute washes of 1x TBST, followed by a 1-hour room temperature incubation of either 1:5,000 in 3% milk α-rabbit or *α*-mouse secondary antibody conjugated to HRP, followed by 3 additional TBST washes. 5-minute activation of HRP was done by development of ECL Clarity Max reagents (Invitrogen). Blots were imaged by chemiluminescence (Biorad ChemiDoc). Each protein of interest was normalized to the band intensity of TATA-binding protein (TBP) using densitometry (ImageLab), and further expression analysis of shFXN normalized to that of the EVEC control.

### CellRox oxidative stress detection

Confluent hBMVEC were incubated with 0.5µM CellRox Deep Red (ThermoFisher) and 0.7µg/ml Hoechst-33324 for 30 minutes at 37°C. Per manufacturer instructions, the cells were washed thrice in PBS before quantifying at 644nm excitation and 665nm emission as normalized to Hoechst fluorescence using the Cytation-Five (Biotek). Representative images are taken at 20x magnification using the Y5 filter cube set of the Cytation-One (Biotek). Representative images are cropped from their original size to emphasize only a few cells per field of view. The size of the scale bar is maintained in the cropping.

### Agilent Seahorse metabolic analysis

Energy metabolism was assessed via Seahorse Mito Stress Test (Agilent). An XF96-well Seahorse plate was coated for five minutes in 0.1mg/ml Poly-D-Lysine, 0.01N HCl, followed by two washes in water and one wash in PBS. Cells were seeded at 8,800 cells per well in a 96-well Seahorse plate in 80µL normal growth RPMI containing serum. 24-hours prior to the assay, the sensor cartridge was equilibrated in sterile water overnight at 37°C without CO_2_, and changed to calibrant for one hour in the same incubation conditions. 36 hours after plating, cell media was removed and replaced with 200µL Agilent assay media (RPMI without phenol red, +10mM glucose, +1mM pyruvate, and +2mM glutamine) for one hour, incubating at 37°C without CO_2_. The media was aspirated and again replaced with 180µL assay media. The sensor cartridge was loaded so that cells were treated at a final concentration of 2.5µM Oligomycin in the A injection port, 1 µM FCCP in the B injection port, and a cocktail of 0.5 µM each Rotenone and Antimycin-A in port C per manufacturer instructions. Each experiment was preceded by the sensor calibration. Following the assay, the wells were washed once in PBS, and lysed using NP-40 based buffer. Protein content as quantified by BCA was used for normalization.

### MitoBright and Mitochondrial Morphology Analysis

hBMVEC were seeded on sterile coverslips and allowed to reach 50% confluence. The protocol was followed per manufacturer instructions. Briefly, cells were treated with mito-bright at a concentration of 1:1,000 with 0.7µg/ml Hoechst-33342 in culture media for 15 minutes at 37°C. The coverslips were washed twice in PBS before fixing for 10 minutes in 3.7% paraformaldehyde, 4% sucrose in PBS at room temperature. The coverslips were washed twice again before mounting to glass slides with prolong-gold antifade mounting media (ThermoFisher). Images were acquired using the 63x oil immersion objective of the Leica DMi8 inverted microscope. At least 6 images were taken per coverslip for three individual experiments. The mitochondrial network of individual cells was analyzed using the Mitochondrial Morphology macro on ImageJ (NIH, Bethesda, MD) per author instructions ^29^.

### Phalloidin-Texas Red

hBMVEC were seeded on sterile coverslips and allowed to reach 50% confluence before fixation in 3.7% paraformaldehyde, 4% sucrose for 10 minutes at room temperature, followed by blocking in 0.1% BSA and 0.01% Tween-20 for 1 hour at room temperature. Phalloidin-Texas Red (ThermoFisher) was diluted at 1:400 in PBS per manufacturer instructions along with 0.7ug/ml Hoechst-33342 for 45 minutes at room temperature. The coverslips were then washed thrice in PBS and mounted as previously described. Images were acquired using the 63x oil immersion objective of the Leica DMi8 inverted microscope. Total polymerized actin was quantified as the integrated density value of phalloidin divided by that of the Hoechst channel, then normalized to EVEC controls. Peripheral actin was quantified by drawing a line of 0.875 inches (6.405µm) across cell membranes in ImageJ. 2-5 membrane regions of interest were quantified per cell per image and aligned as an XY graph in GraphPad prism as previously described ^30–32^. Along the line at which the membrane starts is considered the “peak value,” and then the neighboring 300nm was assigned as the cortical actin region ^33^. Representative images are cropped from their original size to emphasize only a few cells per field of view. The size of the scale bar is maintained in the cropping.

### Transendothelial Electrical Resistance (TEER) Measurements

hBMVEC were seeded apically in 1µm 24-well transwell inserts (Greiner BioOne) with growth media in both chambers, then polarized with serum-free RPMI containing 300nM sodium selenite and 5µg/ml insulin in the basal compartment 8-hours post plating. TEER measurements were acquired using the Endohm-6G (World Precision Instruments) connected to the MillCell-ERS2 voltammeter (Millipore). Ohms values collected at 24, 48, 72 and 96-hours post-plating were subtracted from an identical blank transwell with no cells. Blank-corrected ohms values were then multiplied by the growth area of the 24-well transwells (3.36cm^2^). At 96-hours post plating, apical media aliquots were removed and assayed for lactate dehydrogenase (LDH) secretion (Cytox-96, ThermoFisher) to measure cell death. To account for any seeding differences, the cell monolayer was lysed in 200µL of NP-40 and quantified for protein content using the BCA method for normalization.

### Paracellular tracer flux

hBMVEC were seeded apically in 1µm 24-well transwell inserts (Greiner BioOne) with growth media in both chambers, then polarized with serum-free RPMI containing 300nM sodium selenite and 5µg/ml insulin in the basal compartment 8-hours post plating. At 24-hours in culture, (16-hours post-polarization), the transwells were incubated apically with growth media containing 50µM Lucifer Yellow (LY). Basal media aliquots were removed at 8, 24, 48, and 72-hours post-incubation with LY, quantified for fluorescence at 428nm excitation, 536nm emission, and calculated against a standard curve to generate moles fluxed. At 72-hours post-incubation with LY, apical media aliquots were removed and assayed for lactate dehydrogenase (LDH) secretion (Cytox-96, ThermoFisher) to measure cell death. To account for any seeding differences, the cell monolayer was lysed in 200µL of NP-40 and quantified for protein content using the BCA method for normalization.

### Proliferation Assay

Proliferation kinetics were quantified using CyQuant Red per manufacturer’s instructions. EVEC and shFXN hBMVEC were seeded at 5,000 cells per well of a 96-well plate and left to adhere for 24 hours. At 24-, 48-, 72-, and 96-hours post-plating, cells were incubated with the nuclear dye and background suppressor in growth media per manufacturer instructions for 1hr at 37°C. Cell number as represented by RFU was quantified at 622nm excitation and 645nm emission, respectively on the Cytation5 (Biotek). Cell proliferation as a function of time was calculated by normalizing each day to the “starting” concentration of the 24hr timepoint. A linear regression equation was fit to the growth curve to determine the kinetics of cell growth of each line, and each slope compared.

## Statistics

Graphpad prism was used for generation of all graphs and corresponding statistical analyses. Datapoints were considered outliers if falling outside of the range of two standard deviations from the mean and excluded from analysis. All datasets comparing *E*mpty *V*ector *E*ndothelial *C*ell (EVEC) control and shFXN were analyzed by Student’s T-test, all with a confidence interval at 95%. The F-test was performed to test if the sample groups had significantly different standard deviations. If true, the Welch’s correction was used, also at a confidence interval of 95%. FMS has full access and claims full responsibility for integrity of the data acquisition and analysis. Full data are available upon reasonable request.

## Results

### FXN is knocked down via shRNA in hBMVEC

FXN is an essential protein, and therefore FXN-*knockout* cell and animal systems are non-viable. Thus, we have chosen lentiviral delivery of shRNA as it allows for stable and long-term transfection resulting in a cell line that has a fractional level of remaining FXN within the range found clinically in FRDA patients. In addition, the use of a non-clonal shRNA cell line is advantageous as it provides data from a heterogeneous population, which is true of different cell types in a given patient, and of the patients in a cohort. Indeed, many *in vitro* FRDA studies use similar methods of knock*down* to avoid the lethality of a knock*out* system *e.g.* ∼60% FXN protein expression was observed in a shFXN neuronal model ^34–37^.

Neither the presence nor the role of FXN have been investigated in hBMVEC. The lentiviral-mediated shRNA targeting FXN used in our *in vitro* knockdown model was compared to the empty-vector backbone control plasmid (Horizon Discoveries). Each shRNA plasmid was packaged into the second-generation lentiviral enveloping systems PMD2.G and psPAX2 in HEK293T cells, and viral particles were transfected into Wild-Type Endothelial Cells (WTEC) using polybrene. All experiments using the shFXN hBMVEC are compared against the Empty Vector Endothelial Cell (EVEC) control. Integration of the shRNA was confirmed by GFP expression. (Supplemental Figure 1).

We quantified FXN knockdown using Real-Time quantitative Polymerase Chain Reaction (RT-qPCR) amplifying the FXN transcript in shFXN and EVEC lines. Beta-2-Microglobulin (B2M), a subunit of the constitutively expressed cell surface component MHC class II was used as the housekeeping reference gene to avoid using the classical GAPDH (metabolic enzyme) or *β*-actin (cytoskeletal protein) that would likely reflect changes in energy metabolism and cytoskeletal physiology. The shFXN hBMVEC used in this work retained only 46% of the FXN transcript of the EVEC controls (Fig. 1A).

**Figure 1.**
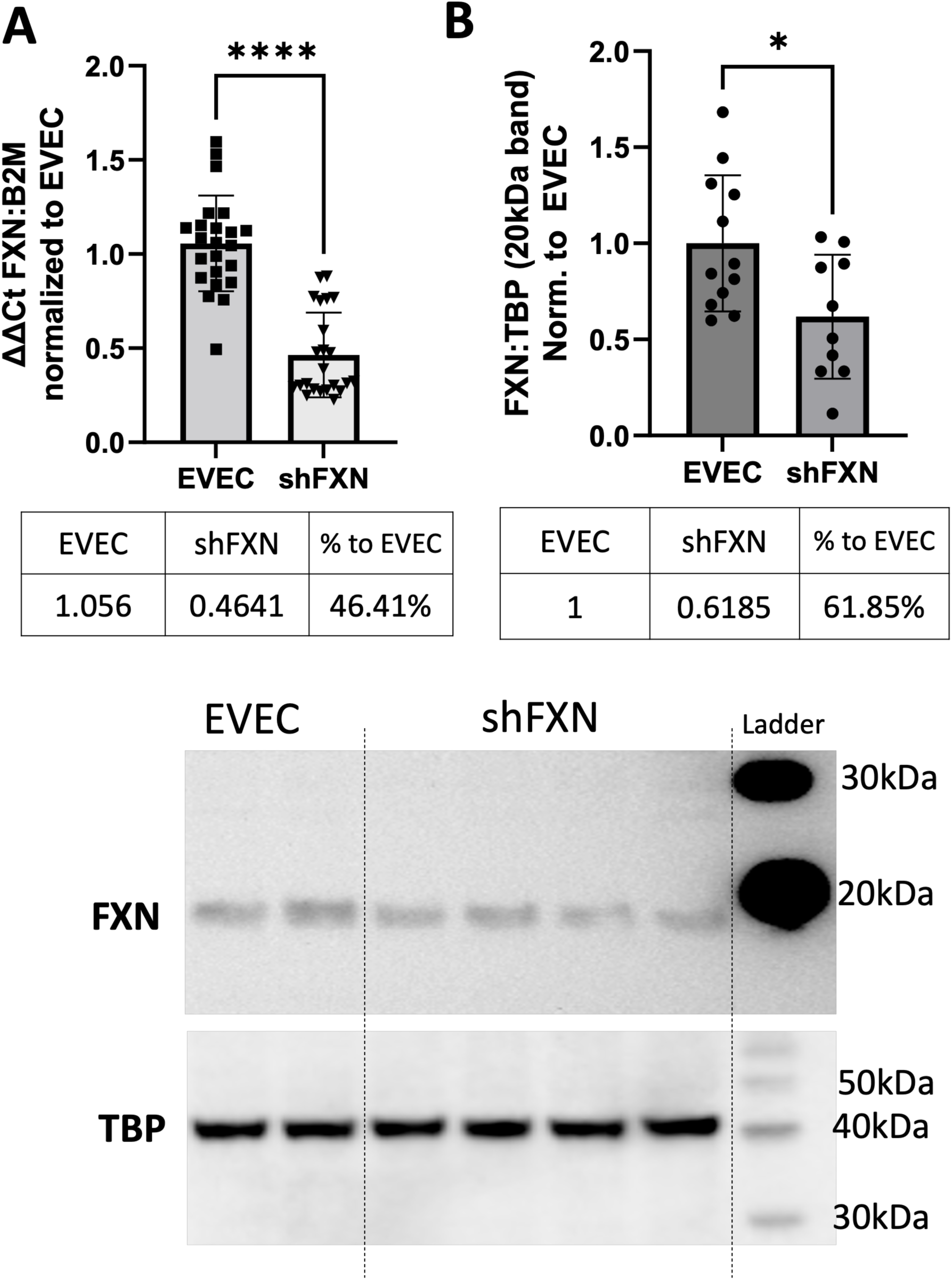
Lentiviral-mediated short-hairpin RNA (shRNA) sufficiently knocks-down frataxin (FXN). (A) FXN transcript is significantly decreased in shFXN. RNA was reverse transcribed and amplified for FXN and Beta-2-microglobulin (B2M) as a housekeeping control, transcript abundance was quantified using the ΔΔCt method normalized to the empty vector endothelial cells (EVEC). (B) Total protein lysates were run on a bis-tris SDS-PAGE, transferred to nitrocellulose, and probed for FXN and Tata-Binding Protein (TBP) as a housekeeping gene. FXN protein expression is normalized to that of TBP as quantified by densitometry, represented as normalized to pooled EVEC values. Representative blot is shown below. Student’s T-test α = 0.05. * p < 0.05, **** p <0.0001. EVEC and shFXN; n= 23. (B) EVEC; n = 12, and shFXN2; n = 10.

We used western blotting to determine if FXN protein levels were decreased as well. EVEC and shFXN lysates were electrophoresed on a 12% bis-tris gel and transferred to nitrocellulose to probe for FXN and the housekeeping control TATA-binding protein (TBP). TATA-binding protein is a transcription factor not considered to be affected by FXN, and therefore used for normalization ^38, 39^. Consistent with transcript abundance, shFXN hBMVEC retain 61% of normal FXN levels present in the EVEC control (Fig. 1B). Our blots represent and quantify the intermediate form of the FXN protein, running ∼18 kDa ^40, 41^. The use of this 18 kDa band was validated using a FXN-overexpression HEK293T lysate, in which the strongest band is shown at ∼18 kDa (Supp. Fig. 3). The FXN precursor form is cleaved twice prior to its mitochondrial translocation, however, neither the precursor (23 kDa) nor the mature (13 kDa) forms of the protein were visible in our western blots ^42^. In summary, our lentiviral-mediated shRNA knockdown of FXN in hBMVEC resulted in 46% transcript and 61% protein remaining in our FRDA hBMVEC model (Fig. 1D).

### shFXN hBMVEC exhibit increased oxidative stress

Oxidative stress is a known hallmark of FXN loss; this stress is downstream of reactive oxygen species (ROS) formed by free, non-chaperoned iron ^12, 43^. CellRox Deep Red was used as an indicator of intracellular oxidative stress in the shFXN and EVEC cell lines. CellRox is reactive with all forms of ROS. EVEC (Fig. 2A) and shFXN (Fig. 2B) treated with 0.5 μM CellRox were analyzed for total intracellular oxidative stress spectrofluorometrically as normalized to Hoechst. Indeed, shFXN hBMVEC exhibited increased intracellular oxidative stress (+8.9%) as compared to EVEC (Fig. 2C). Increased oxidative stress in this model is consistent with decreased FXN, and correlates with the oxidative modification of actin glutathionylation that is observed in FRDA patient fibroblasts ^17, 24^.

**Figure 2.**
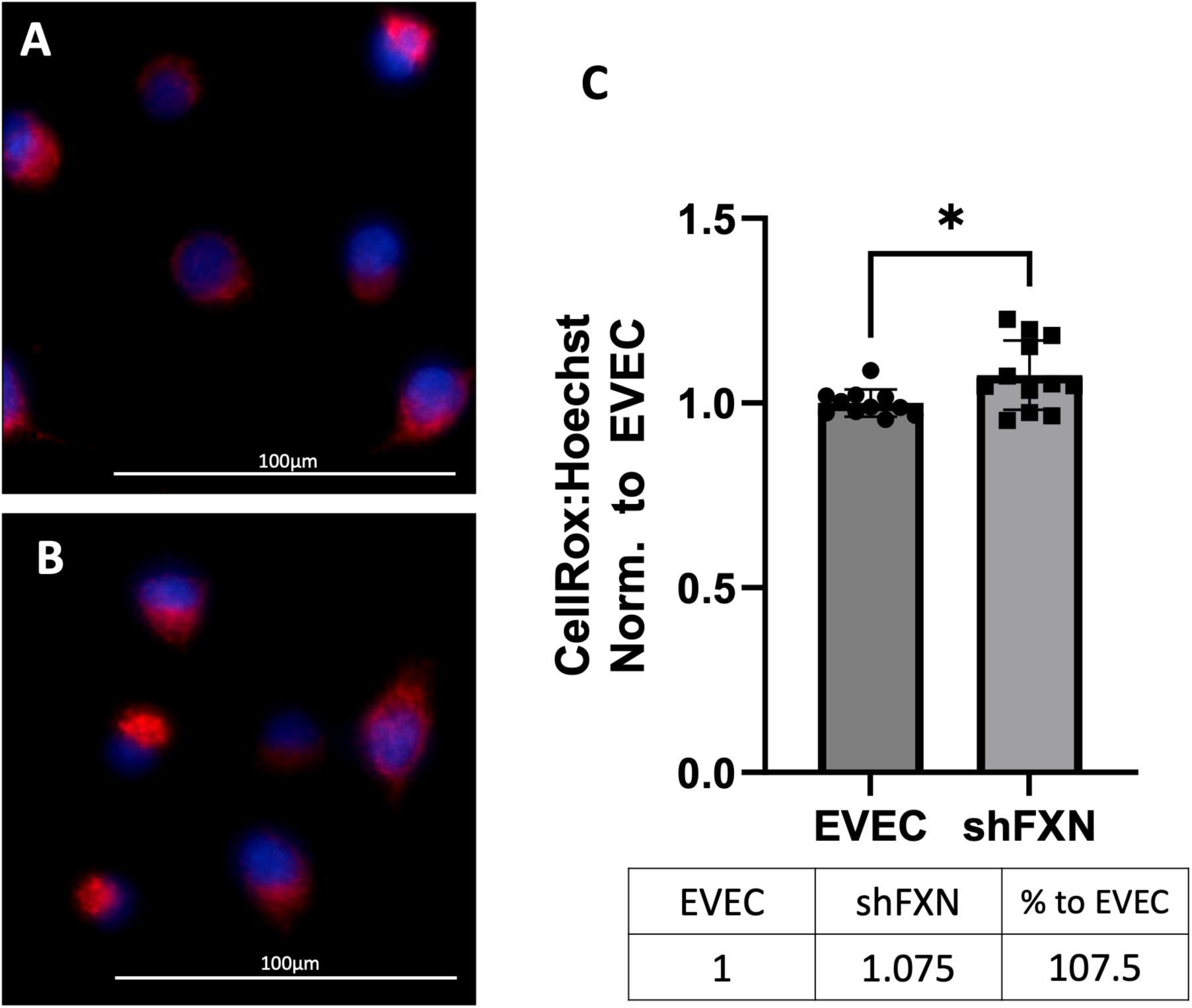
shFXN have upregulated oxidative stress. (A) EVEC and (B) shFXN are incubated with 0.5µM CellRox Deep Red and 0.7µg/ml Hoechst for 30 minutes at 37°C. (C) Fluorescence values are read on the Cytation-5 (Biotek), the Cellrox normalized to Hoechst values, and then normalized to EVEC values to represent a fold change. Student’s T-test, α=0.05. *; *p*<0.05. EVEC and shFXN n=8. Representative of 3 biological replicates. The images in panels A and B were cropped from their original size while maintaining the scale of the scale bar.

### shFXN hBMVEC have decreased oxidative energy metabolism and ATP production with shift to glycolysis

Another hallmark of FXN loss is decreased metabolic capacity due to lack of iron incorporation into the electron transport chain (ETC) complexes I-III and aconitase,^7, 8, 44, 45^. The loss of aconitase caused by FXN deficiency likely reduces Krebs cycle generation of ETC-requiring reducing equivalents ^44^. In addition, decreased activity of the ETC would reduce production of the NAD+ required for glycolytic ATP production, creating a negative feedback loop with respect to energy production overall. Thus, the reduction in ETC activity in FRDA can be expected to directly reduce oxidative production of ATP and secondarily depress glycolytic energy metabolism.

We have measured both OXPHOS and glycolysis in our model using the Agilent Seahorse Mito Stress Test. Indeed, shFXN hBMVEC are significantly deficient in oxygen-mediated energy metabolism compared to the EVEC controls as represented by nearly 30% reduction in the oxygen consumption rate (OCR) (Fig. 3A). We were intrigued to see that the shFXN hBMVEC has a slight, albeit non-significant, decreased marker of glycolytic function (extracellular acidification rate (ECAR)) (Fig. 3B). However, the exact influence of FXN knockdown on glycolysis cannot be known as ECAR is representative of not only glycolytic function, but non-mitochondrial respiration as well, as it is influenced by CO_2_ production from the Kreb’s cycle ^46^.

**Figure 3.**
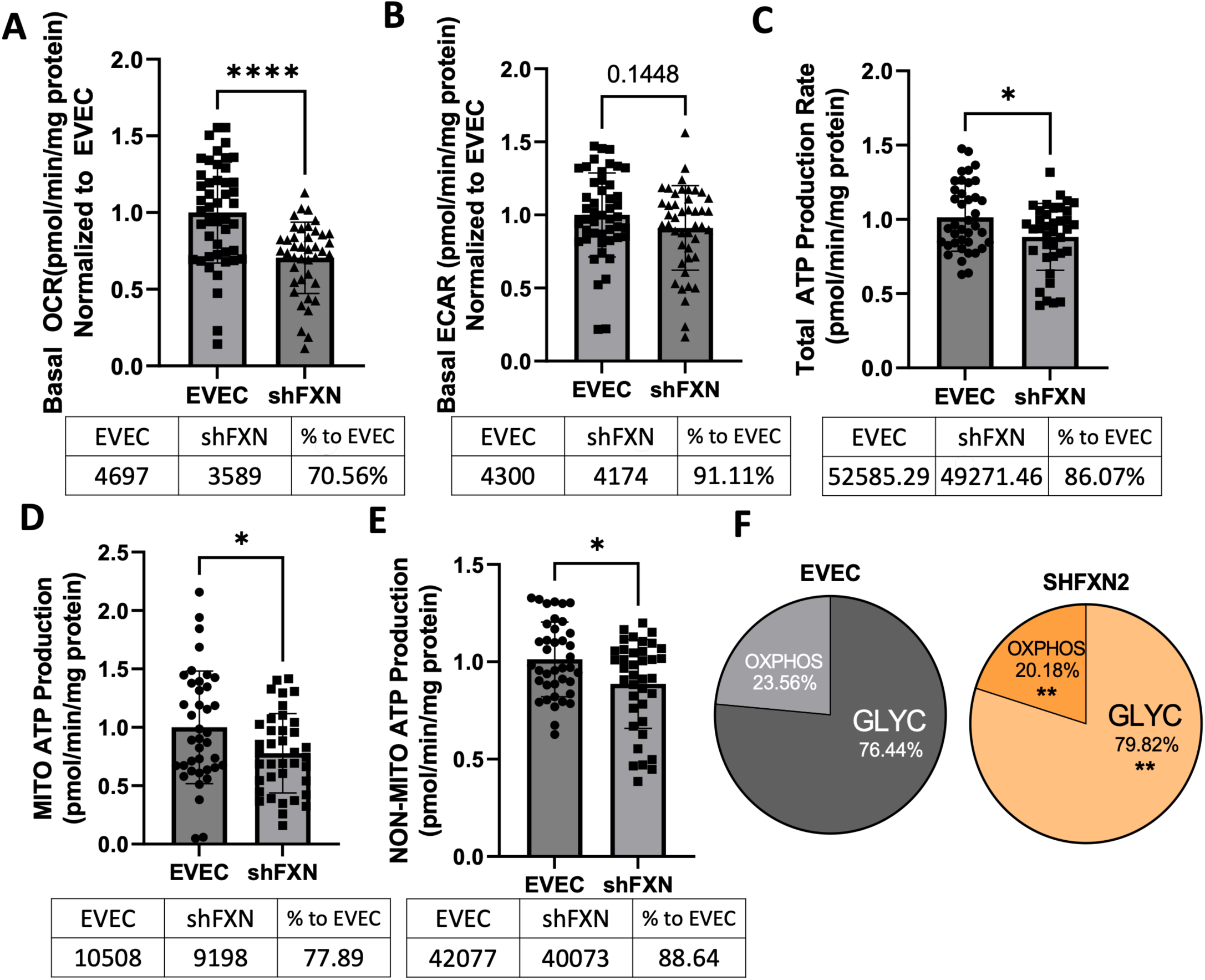
Metabolism is affected by Frataxin-knockdown in hBMVECs. Basal levels of (A) Oxygen consumption rate (OCR), (B) extracellular acidification rate (ECAR), (C) Total ATP production, (D) ATP produced via mitochondria, and (E) ATP produced by glycolysis are measured in the Agilent Seahorse Mito Stress Test and then normalized to mg protein in the well. (F) The percentage of oxidative phosphorylation and glycolysis. Student’s T-test α =0.05; * *p* < 0.05, ** *p* < 0.01, **** *p* <0.0001. (A). EVEC; n = 47, shFXN; n = 41. (B) EVEC; n = 47, and shFXN; n = 44. (C, D, E and F) EVEC; n = 39, shFXN; n = 38. Representative of 2 biological replicates.

In total, the shFXN hBMVEC are deficit in energy production, with total ATP levels decreased by 14% compared to EVEC controls (Fig. 3C). In addition, the levels of ATP produced by OXPHOS (Fig. 3D) and glycolysis (Fig. 3E) were significantly decreased in shFXN hBMVEC, by ∼12% and 11%, respectively. Additionally, shFXN hBMVEC are shifted to an increasingly glycolytic phenotype compared to the EVEC controls (Fig. 3F), likely due to the direct role of FXN in ETC function, causing decreased NAD+ availability as a secondary effect on glycolytic function. Furthermore, modest decrease in shFXN ECAR (Fig. 3B) *versus* significant decrease of non-mitochondrial ATP production (Fig. 3E) might indicate uncoupling of ATP production in the glycolytic pathway, as three hydrogen atoms are produced before the final ATP hydrolysis during pyruvate production ^47^. These metabolic markers align with the known OXPHOS pathologies in FRDA models, as well as revealing a related deficiency in glycolytic ATP production as well.

### shFXN have more mitochondrial objects and larger cell size

Due to FXN’s role in mediating mitochondrial physiology, we sought to determine the effects of FXN knockdown on mitochondrial networking. Aberrant mitochondrial dynamics in both fission and fusion pathways have been reported in FRDA ^48^. To investigate mitochondrial networking in our shFXN hBMVEC, we used Mitobright Deep Red to stain mitochondria, and the Mitochondrial Morphology ImageJ Macro for network analysis ^29^.

shFXN showed increased levels of mitochondrial number consistent with mitochondrial fission defects in FRDA patient-derived cardiac iPSCs ^48^ (Fig. 4A). We were surprised to see that the cytosolic occupancy of mitochondria, however, was nearly the same across the cell samples (Fig. 4B). Cytosolic occupancy is a density ratio of mitochondrial objects normalized to cell size; and so we were interested in analyzing cell area. Indeed, shFXN hBMVEC are significantly larger than EVEC controls, representing a 17% increase in cell area (Fig. 4C). FRDA patient fibroblasts with increased actin glutathionylation also had increased cytosolic complexity revealed by side-scatter flow cytometric analysis and increased cell size ^17, 24^.Therefore, the observed changes in shFXN cell size led us to investigate potential changes in actin filament abundance and distribution.

**Figure 4.**
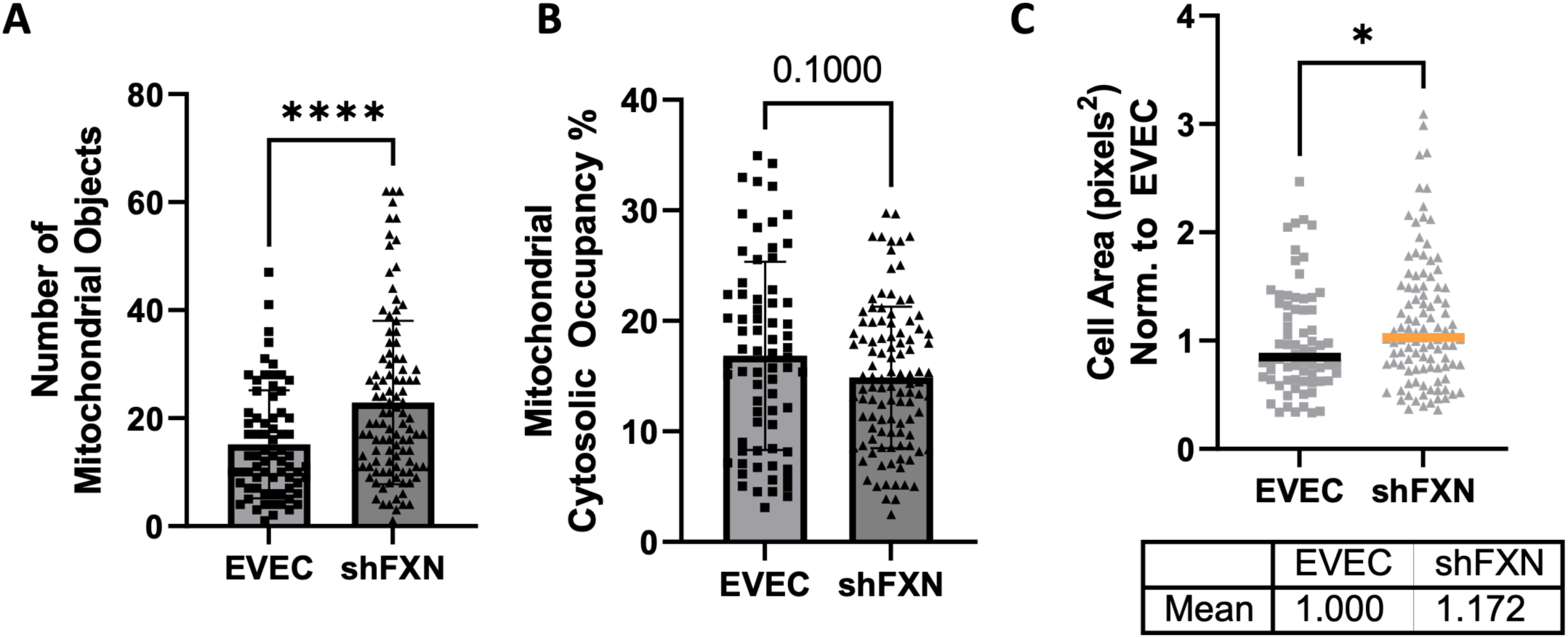
Mitochondrial physiology is altered in shFXN. (A) Mitobright mitochondrial dye is used to determine the number of mitochondrial objects per cell. (B) Mitochondrial number is normalized to the cell area to determine the percentage cytosolic occupancy. (C) Total cell size is quantified and normalized to EVEC controls. Student’s T-test α =0.05 * p < 0.05, **** p < 0.0001. (A) EVEC; n = 71, shFXN; n = 105. (C) EVEC; n = 71, shFXN; n = 106 (B). EVEC; n = 71, shFXN; n = 105. Each representative of 3 biological replicates.

### shFXN have decreased levels of polymerized total and peripheral actin

Early reports of FRDA cell biology cited larger cell size along with increased actin glutathionylation in patient-isolated fibroblasts ^17^. Research has shown that actin glutathionylation inhibits actin polymerization dynamics downstream of oxidative stress, decreasing both the rate of filament polymerization and filament length ^18^. Filamentous actin (F-actin) is essential in barrier physiology due to its anchorage of tight junction proteins *via* the scaffolding protein ZO-1, and outside of the membranous actin sits the cortical actin ring (CAR), a network of actin fibers that in addition to providing junctional support also increases structural integrity and extracellular matrix adhesion.

Using phalloidin-Texas red, a dye which only stains F-actin, we quantified total, membranous, and CAR actin fibers in shFXN and EVEC lines. We quantified a significant loss of total F-actin in shFXN (Fig. 5B) hBMVEC compared to EVEC (Fig. 5A); this difference represented a nearly 20% F-actin reduction (Fig. 5C).

**Figure 5.**
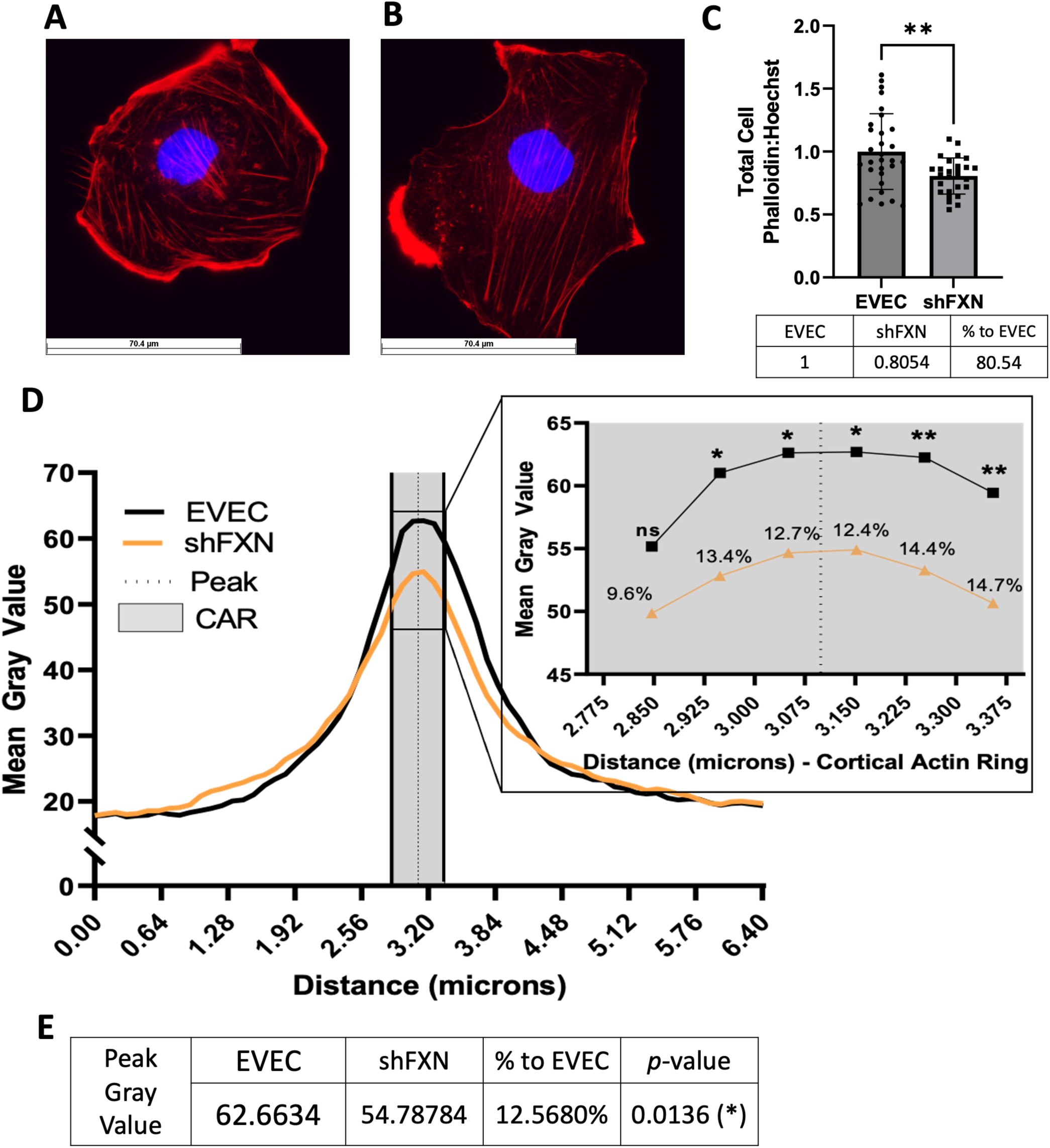
shFXN hBMVEC are deficient in total polymerized actin and lack peripheral actin. (A) EVEC and (B) shFXN hBMVEC are incubated with Phalloidin-Texas Red and Hoechst, (C) the ratio taken to quantify total filamentous actin per cell, then normalized to EVEC controls. (D) 6.4 µm of 2-5 membranes per cell were quantified for F-actin (gray value) and averaged into a histogram. The membrane interface (peak) is represented by the dotted line. 300nm surrounding the peak was quantified as the cortical actin region (CAR), as represented by the shaded region. Individual datapoints of the cortical ROI are represented in the inset with significance and the %deficit in F-actin. (E) Values and %difference with statistics for the peak membrane F-actin. Student’s T-test α =0.05; ns = not significant, * p < 0.05, ** p < 0.01. Total phalloidin analysis (C): EVEC; n = 30, shFXN; n = 27. Membranes analyzed (D,E): EVEC; n = 191, shFXN; n = 222. The images in panels A and B were cropped from their original size while maintaining the scale of the scale bar.

A difference in Phalloidin staining at the cell membrane was apparent, and so we created a histogram of Phalloidin Texas-Red pixel density along a 6.4 μm line drawn through the cell membrane ^31, 32^. The line was drawn so that the membrane interface represented the middle of the line to generate a bell-curve, averaging 2-5 membrane regions of interest per cell per image (Fig. 5D) ^30^. This analysis revealed that the shFXN (orange) trace had notable differences in F-actin distribution compared to the EVEC controls (black). This difference was at a maximum at the center of the curves, which represents F-actin at the membrane interface. The membrane peak is indicated by the dotted line drawn through the curve, which shows that shFXN hBMVEC have 12.5% less F-actin at the cell membrane (Fig. 5E). As mentioned, the CAR is also essential to barrier integrity by providing structural integrity, adhesion, and junctional support. CARs can span 10-300 nm below the membrane surface, and so 300 nm regions flanking the membrane peak are represented in the gray shaded region (Fig. 5D) ^33, 49, 50^. Furthermore, the CAR region is maximized in the inset (Fig. 5D). The shFXN hBMVEC have significantly less F-actin in 5 of the 6 CAR datapoints, ranging from ∼10-15% less than the EVEC controls.

We questioned if the changes in F-actin of shFXN were downstream of deficient *β*-actin transcriptional and translational processing. To assess this, we used qPCR, comparing β-actin to the housekeeping control B2M and found no transcriptional changes between the EVEC and shFXN hBMVEC (Supp. Fig. 3A). Western blot analysis using the house-keeping control TBP also revealed no significant changes in actin abundance. Thus, we observed a significant decrease in filamentous actin in shFXN at the whole-cell level, at the cell membrane, and in the cortical actin ring. This indicates that the cytoskeleton is altered downstream of FXN-loss, and that physiological functions relying on the actin cytoskeleton likely are affected in disease.

### shFXN hBMVEC exhibit increased paracellular permeability

We next determined if the increase in cell size and the loss of polymerized total and cortical actin resulting from FXN knockdown altered hBMVEC barrier integrity. To do so, we plated hBMVEC on mesh transwell membranes which facilitate solute flux between two chambers as a model of cell barrier. hBMVECs were plated on the apical side of the membrane, and then polarized and incubated apically with 50 µM Lucifer Yellow (LY) at 8-hours in culture. At 24, 48, 72, and 96-hours in culture, hBMVEC were assessed for transendothelial electrical resistance (TEER) and LY flux (Fig. 6A). LY is a cell-impermeant fluorescent tracer that is not taken-up or transcellularly trafficked through cells, and so its passage across a transwell system is indicative *only* of paracellular permeability. To account for any discrepancies in initial seeding density, each transwell was lysed at the final timepoint and assessed for protein content, which was used as a normalization factor (Supp. Fig. 4A).

**Figure 6.**
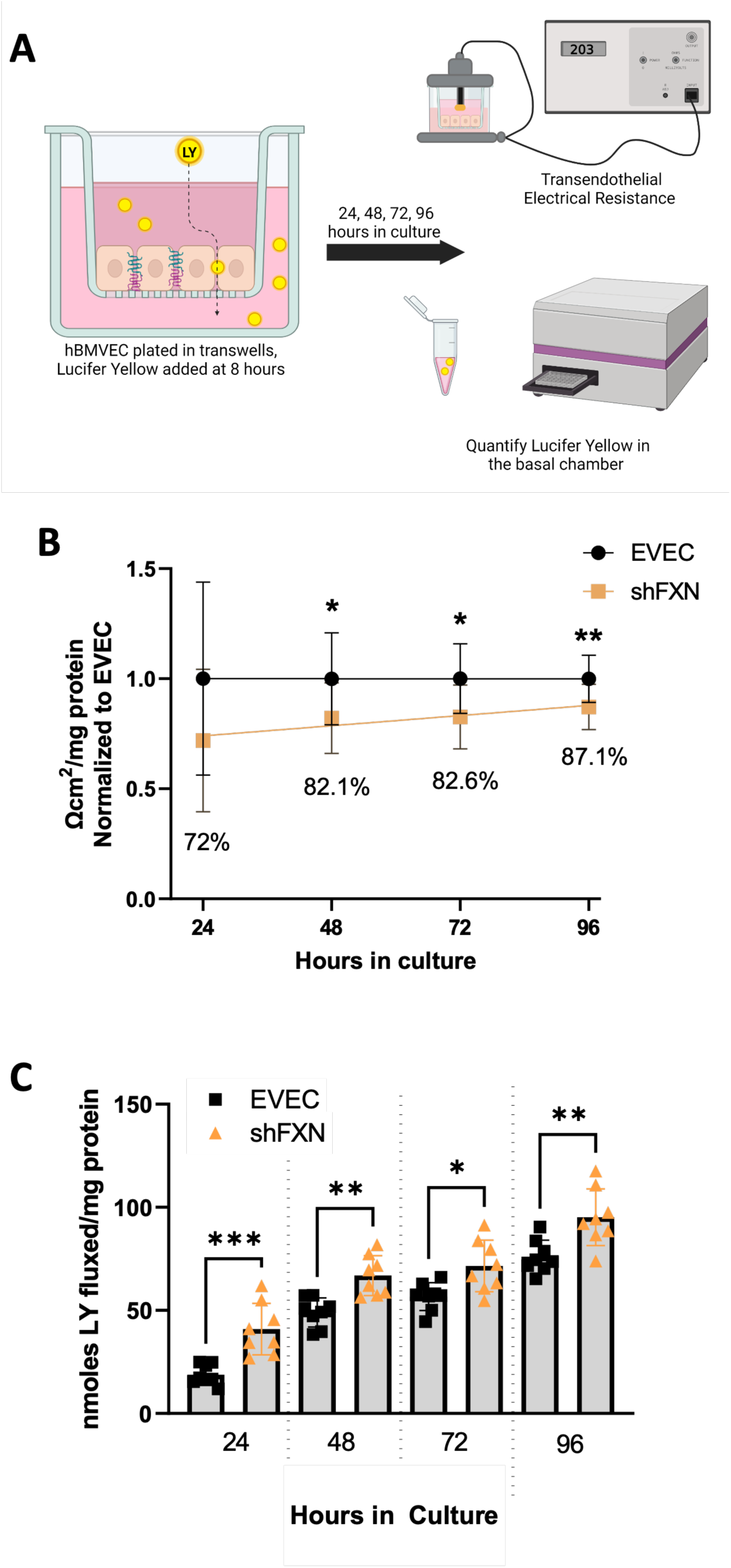
Paracellular permeability is increased in shFXN. (A) hBMVEC plated in the apical chamber of transwells are treated with 50µM Lucifer Yellow (LY). Transendothelial electrical resistance (TEER) measurements and LY aliquots are taken at 24, 48, 72, and 96-hours in culture. (B) TEER is measured using the Endohm-6G (World Precision Instruments), sample wells are background corrected to a transwell not containing cells. Blank-corrected values are multiplied by the growth area of the 24-well transwell (3.36cm2) and normalized to the final protein content of the well. Represented as fold change to EVEC per timepoint. (C) Media from the basal chamber is analyzed for nanomoles of LY fluxed at 24-96 hours in culture at 490nm and 525nm excitation and emission, respectively. The cell monolayers were lysed after the final timepoint, and total protein content was used for normalization of all experiments. Student’s T-test α =0.05; * p < 0.05, ** p < 0.01, *** p < 0.001. (B) EVEC; n =11, and shFXN; n = 12. (C) EVEC; n = 8, and shFXN; n = 7. Representative of 2-4 biological replicates.

At each of the timepoints, there is a clear deficit in the shFXN (orange) barrier integrity as measured by TEER compared to EVEC (black); this deficit ranged between 30% to 13% over this time course (Fig. 6B). While the shFXN cells approach the barrier strength of the EVEC control at later timepoints, they never reach the resistance values of the control barrier. Additionally, increased permeability is apparent with significantly increased amounts of LY flux into across the shFXN barrier relative to EVEC control, each timepoint assayed (Fig. 6C).

“Apparent” permeability could result from any one of the following: 1) true paracellular permeability, 2) cell death creating holes in the cell monolayer, or 3) slower growth kinetics of the shFXN hBMVEC cells. To evaluate increased cell death, we removed apical media aliquots on the final day of the experiment to assay for cell death as indicated by LDH release (Supp. Fig. 4B). The shFXN hBMVEC experienced *no* increase in LDH release compared to EVEC controls. To discount the possibility that altered growth kinetics lead to a deficiency in barrier integrity, we measured culture growth along the same 24, 48, 72, and 96-hour timeline of the permeability experiments using the nuclear dye CYQUANT Red (Supp. Fig. 4C). The 48, 72, and 96-hour timepoints were normalized to the 24-hour timepoint to account for any possible discrepancies in seeding density. A linear regression was fitted to the linear portion of the growth curve showed that the shFXN (orange) and EVEC hBMVEC (black) proliferate at statistically similar rates.

## Discussion

FRDA patients, *in vitro* animal models, and *in vitro* biochemical data are suggestive of BBB breakdown, but this premise has not been explicitly interrogated. Examination of this premise may be of particular relevance with respect to the neuroinflammation, brain iron accumulation, neurodegeneration, and stroke experienced by FRDA patients.

Clinically, 20% of FRDA patients experience stroke ^51–53^. Notably, brain iron accumulation follows neurodegeneration but is progressive, suggesting that brain iron deposition occurs throughout disease ^14^. Similarly, cerebellar sodium content is increased in FRDA patients outside of significant atrophy, something strongly associated with development of edema ^14, 54, 55^.

Molecularly, the inflammatory cytokine IL-6 is upregulated in FRDA circulation, a pathway which in neurodegeneration is linked to decreased protein expression of occludin (TJ protein) and cadherin (adherens junctions), causing BBB breakdown. IL-6 also activates the M1 inflammatory microglial phenotype, which is observed in FRDA, and both contribute to BBB-breakdown and hyperpermeability in other models of neurodegeneration ^56–58^. Further compounding evidence for brain iron accumulation via *paracellular* rather than *transcellular* flux is indirect downregulation by IL-6 of the iron exporter ferroportin ^59–63^.

In line with inflammation, barrier dysfunction is also intimately linked to loss of Nrf2 transcription factor function as seen in brain, intestine, and lung ^64–66^. Related to brain vasculature, siRNA-mediated knockdown of Nrf2 in bEnd.3, a mouse BMVEC line, and inflammatory insult caused by LPS treatment decreased transcriptional profiles of ZO-1 and occludin consistent with increased barrier permeability ^67^.

Furthermore, matrix metalloprotease-9 is upregulated in FRDA patient blood samples and the KIKO FRDA mouse model, potentially disrupting the integrity of the vascular basement membrane ^11, 68^. Cyclooxygenases are increased in several FRDA mouse models and isolated FRDA patient B-lymphocytes, and VEGF is increased in FRDA patient olfactory mucosal mesenchymal stem cells, and both pathways are correlated to downregulation of claudin-5 and occludin, the former being essential to brain vasculature 69-72 73.

The FXN-knockdown hBMVEC model system described here provides a platform to examine the role brain vascular homeostasis plays in the cerebral pathophysiology in FRDA. Our shFXN cell line displays at the cell level the hallmark pathologies of FRDA, including increased oxidative stress (Fig. 2), decreased total energy production (Fig. 3), aberrant mitochondrial networking (Fig. 4A), and increased cell size (Fig. 4C). Our shFXN model may be considered a mild knockdown due to the retention of 61% of protein expression in the shFXN hBMVEC compared to EVEC, whereas classical FRDA patients retain on average only 30%. It should be noted that while the GAA-expansion mutation in teenage years is the most common FRDA cause, ∼2-5% of patients develop disease due to a single-allele point mutation (pFA), and “late onset Friedreich’s Ataxia” (LOFA) represents about 25% of the disease cohort ^74, 75^. Residual FXN protein levels vary in these cases, with 33% FXN retention in pFA, and 65.6% in LOFA ^75^. Therefore, while our model has a mild knockdown compared to classical FRDA and pFA, it is within the range of FXN knockdown linked to disease, and, most critically, displays FRDA cell pathologies nonetheless. As noted, are hBMVEC model is similar to ∼60% FXN protein expression in shFXN neurons ^37^.

Our model exhibits an interesting energy metabolic phenotype, with a significant loss of oxidative phosphorylation (Fig. 3A) but a marginal decrease in non-oxidative metabolism (Fig. 3B) We hypothesize that FXN knockdown directly decreases ETC-based production of NAD+, thereby diminishing the pool of reducing cofactors required for glycolysis and the Krebs cycle. Consistent with reduction of ETC flux, and the apparent indirect effect on glycolysis, we see the metabolic profile in these cells being shifted more towards glycolysis (Fig. 3F).

Increased cell size is known in FRDA fibroblasts downstream of actin glutathionylation, which we also observe in our shFXN hBMVEC (Fig. 4C) ^17^. Not only is total F-actin decreased in shFXN hBMVEC (Fig. 5C), F-actin presentation at the membrane peak is decreased and at the cortical actin ring surrounding the membrane (Fig 5D-E). Cortical actin is essential in maintaining ZO1, the scaffolding protein of the tight junction family, to the transmembrane domains of Claudin-5 and Occludin tight junction effectors, as well as providing structural integrity and adhesion. Note that the actin deficiency in these cells is not due to transcriptional (Supp. Fig. 3A) or translational (Supp. Fig. 3B) defects of *β*-actin processing, and thus appears post-translationally regulated ^17, 18, 24^.

Based on the changes in cell size and altered actin dynamics in shFXN, we were interested in the paracellular integrity of shFXN hBMVEC compared to the EVEC controls. Indeed, shFXN hBMVEC have significantly decreased barrier integrity, which approaches, but never meets the strength of EVEC control barriers (Fig. 6B and C). Importantly, this increase in permeability was not due to differences in seeding density, increased cell death, or decreased growth kinetics (Supp. 4A-C), suggesting true paracellular permeability downstream of FXN loss. Therefore, we demonstrated our shFXN hBMVEC model has increased paracellular permeability co-incident with increased cell size and loss of total and membranous filamentous actin. These findings contribute to our model that oxidative modifications of actin depress filament formation and junctional integrity, allowing for paracellular solute permeability (Fig. 7).

**Figure 7.**
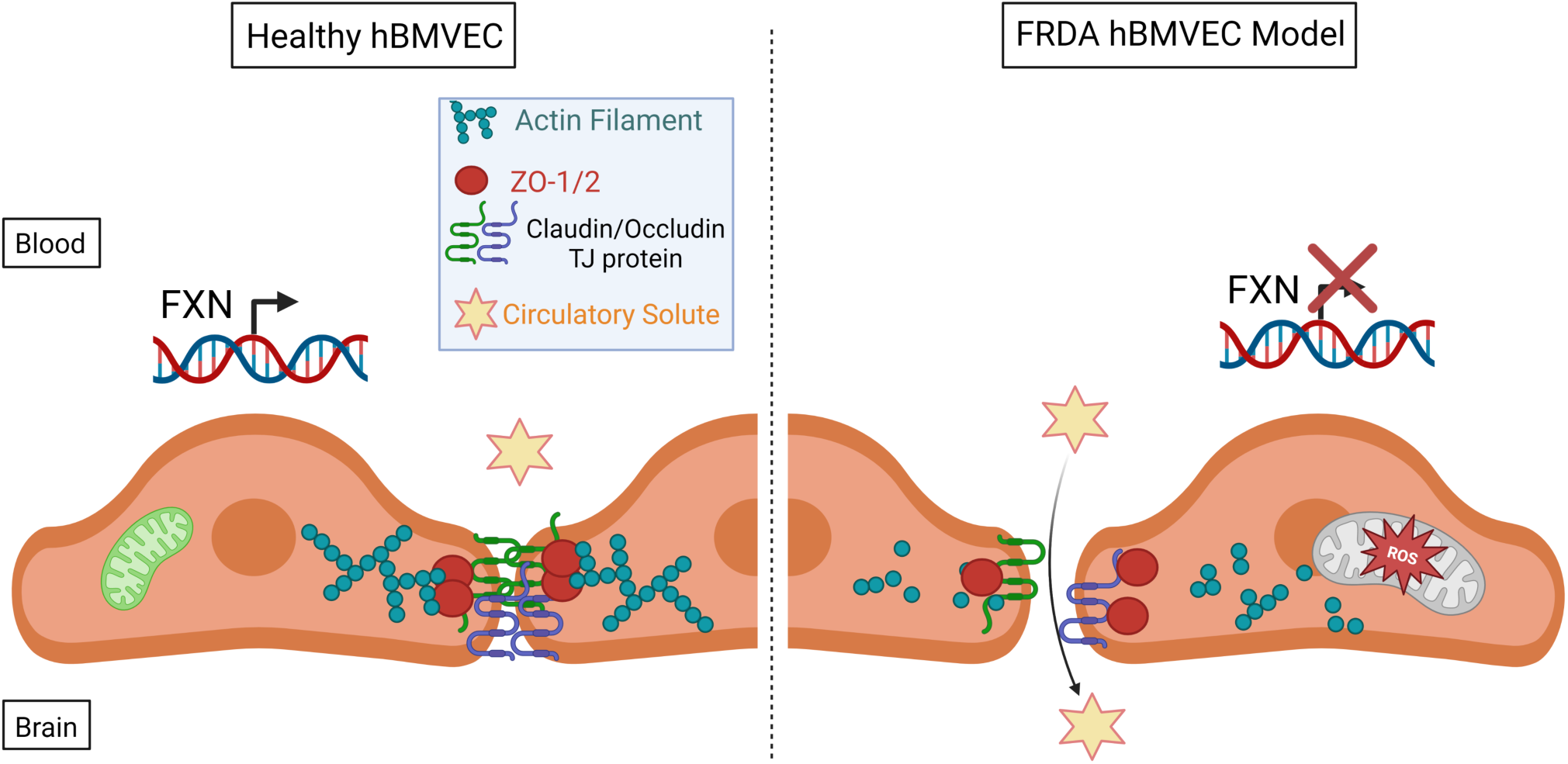
The proposed schematic of FXN-mediated hBMVEC permeability. (A) Healthy hBMVEC maintain FXN processing, have healthy mitochondrial dynamics, and proper actin polymerization to maintain paracellular tight junctions that prevent paracellular flux. (B) FRDA hBMVECs lose FXN transcriptional processing, lack healthy mitochondrial physiology, and have less polymerized actin at the membrane, leading to unsupported tight junctions and paracellular permeability.

While our method of modeling FRDA is shRNA-mediated FXN knockdown, we did not interrogate a *direct* cytoskeletal alteration arising from cis-silencing of *PIP5K1β* via *FXN* expansion tracts ^19^. However, we do see a significant phenotype associated with FXN loss alone, indicating that FXN loss is *sufficient*, though potentially not *fully* responsible for cytoskeletal and barrier alterations in FRDA. Overall, our data indicate loss of blood-brain barrier integrity with FXN loss *in vitro,* thus supporting the examination of the microvasculature, in FRDA and other neurodegenerative disorders as it contributes to the associated organ pathophysiology.

## Conclusions

There are gross anatomy findings and molecular biochemical data which suggest vascular dysfunction in FRDA patients. We have identified significant barrier deficiency in our model of shFXN hBMVEC. These cells display a loss of total filamentous actin, and importantly, significant deficiency of F-actin at the cell membrane and in the cortical actin ring regions. This is consistent with increased barrier permeability and increased paracellular solute flux. These findings provide new insight into brain vascular manifestations of FRDA previously uncharted in studies of FRDA brain pathology. Our data provide a new understanding of BBB function in FRDA, identifying a potential therapeutic target in the neuroinflammation, neurodegeneration, brain iron accumulation, and stroke in this disease.

## Novelty and Significance

### What is known

- FRDA patients experience longitudinal brain iron accumulation that parallels the decline of patient quality of life.
- 20% of FRDA patients suffer a stroke, and stroke causes 7% of FRDA deaths.
- There are both direct (PIP5K1β) and indirect (actin glutathionylation) effects on the cytoskeleton in FRDA.

### What new information does this article contribute?

- shFXN hBMVEC display: increased oxidative stress, loss of energy metabolism, mitochondrial abnormalities, and increased cell size consistent with disease and other FRDA models
- shFXN hBMVEC have loss of filamentous actin at the: whole cell level, at the plasma membrane, and at the cortical actin ring
- shFXN hBMVEC are paracellularly permeable to tracer solutes

Our model of frataxin-deficient human brain microvascular endothelial cells (hBMVEC) is novel in the FRDA literature. While some more recent FRDA reports investigate heart and lung *macro*vascular pathology, consideration of the brain microcapillaries has been overlooked, despite the prevalence of stroke and stroke-related complications in this disease. Many reports connect inflammatory and oxidative signatures to vascular barrier dysfunction, but this has *not* been considered with respect to the molecular bases of FRDA pathology. Our investigation of brain vasculature pathology identifies loss of barrier integrity consistent with cytoskeletal abnormalities, something briefly touched on in early FRDA literature with the identification of actin glutathionylation. Here, we demonstrate that actin filament polymerization in FXN-knockdown hBMVEC is deficient, especially at the plasma membrane and at the cortical actin region, components of the cytoskeleton essential in supporting tight and adherens junctions, cell rigidity, and extracellular matrix adhesion. Consequently, we see an increase in paracellular permeability, in total indicating barrier defects downstream of FXN loss. Our work provides insight on vasculature of the FRDA brain, which could become a clinical drug target in maintaining brain homeostasis, in the context of brain iron accumulation, neuroinflammation, and stroke.

## Acknowledgements

FMS and DJK were involved in conception and design. FMS performed data acquisition and figure formation. FMS and DJK were involved in analysis and interpretation of data. FMS and DJK were involved in preparation, revision, and submission of the manuscript. The authors acknowledge Biorender.com, for which schematic figures were made.

## Supplemental Materials

**Supplemental Figure 1.**
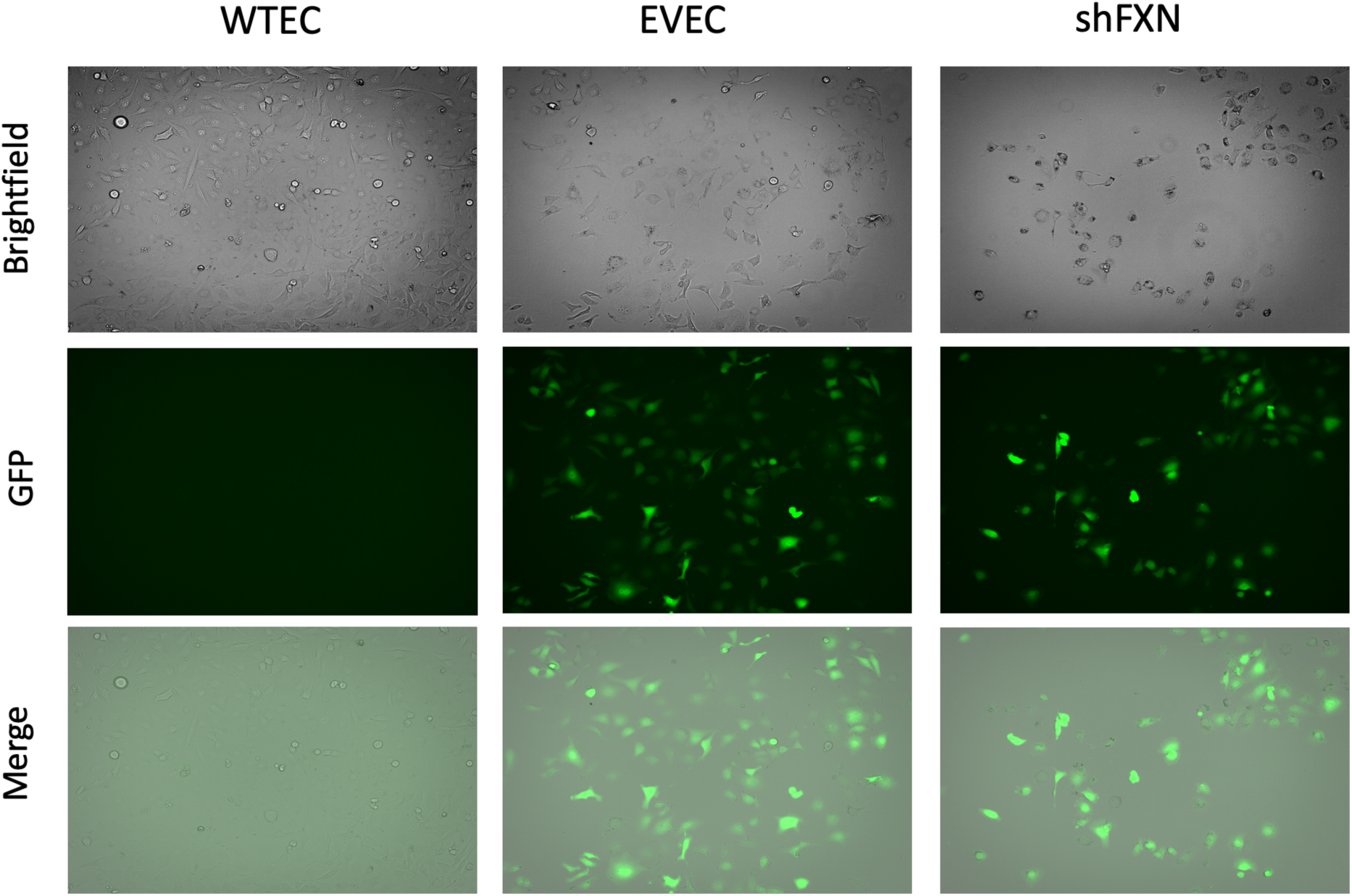
GFP expression confirms integration of lentivirus, which is absent in non-transfected Wild-Type Endothelial Cells (WTEC) but present in lentiviral-transfected Empty Vector Endothelial Cells (EVEC) and shFXN.

**Supplemental Figure 2.**
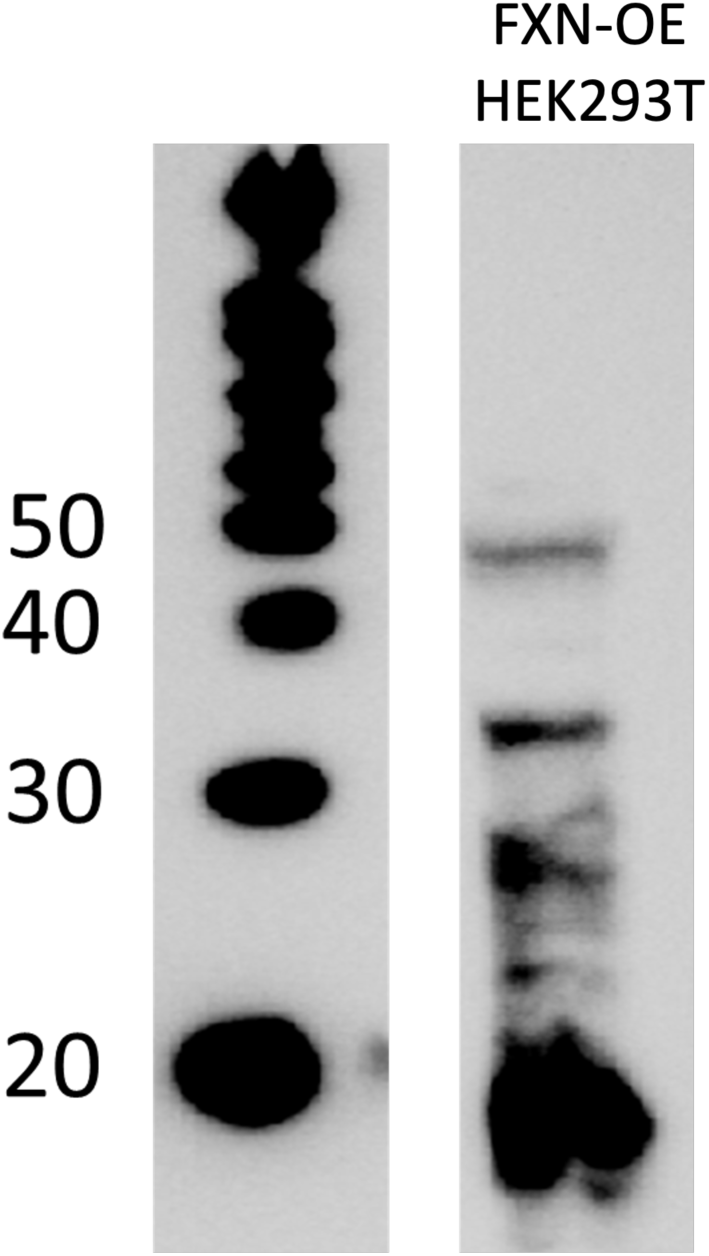
20µg FXN-overexpression HEK293T lysate is electrophoresed and probed for FXN for validation of bands evident with α-FXN primary antibody (ThermoFisher PA5-13411). The ∼18kDa major band corresponds to the mature form of FXN and is therefore used for protein expression analysis in our experiments.

**Supplemental Figure 3.**
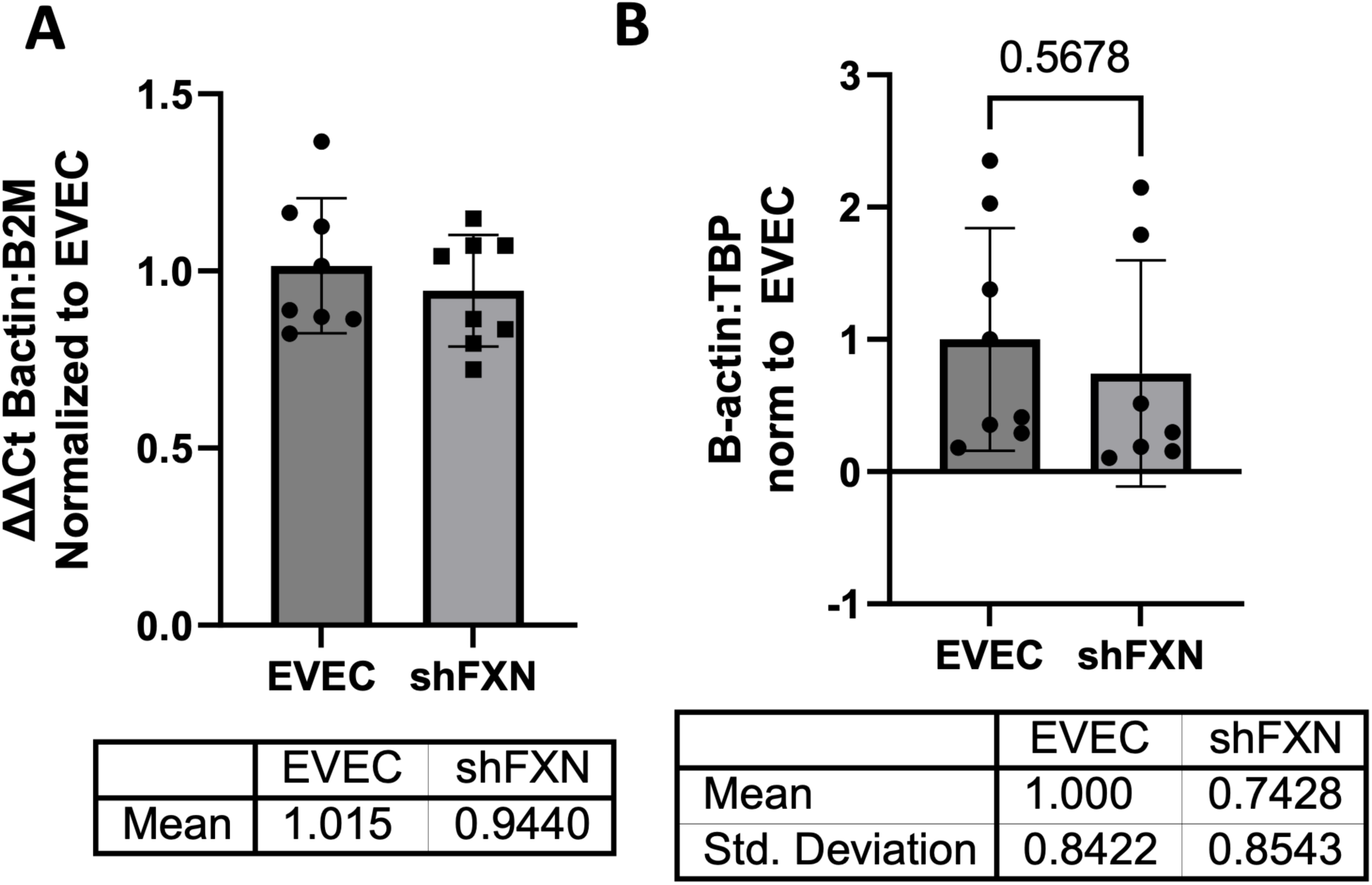
Loss of F-actin in shFXN hBMVEC is not due to transcriptional or translational defects. (A) hBMVEC RNA was reverse transcribed and amplified for β-actin and Beta-2-microglobulin (B2M) as a housekeeping control, transcript abundance was quantified using the ΔΔCt method normalized to the empty vector endothelial cells (EVECs). (B) Total protein lysates were run on a bis-tris SDS-PAGE, transferred to nitrocellulose, and probed for α-β -actin and Tata-Binding Protein (TBP) as a housekeeping gene. β-actin protein expression is normalized to that of TBP as quantified by densitometry. The graph represents the fold-change of β-actin expression in shFXN relative to EVEC controls. (A) EVEC and shFXN; n= 8. (B) EVEC; n=8, shFXN; n=7. Each representative of 2 biological replicates.

**Supplemental Figure 4.**
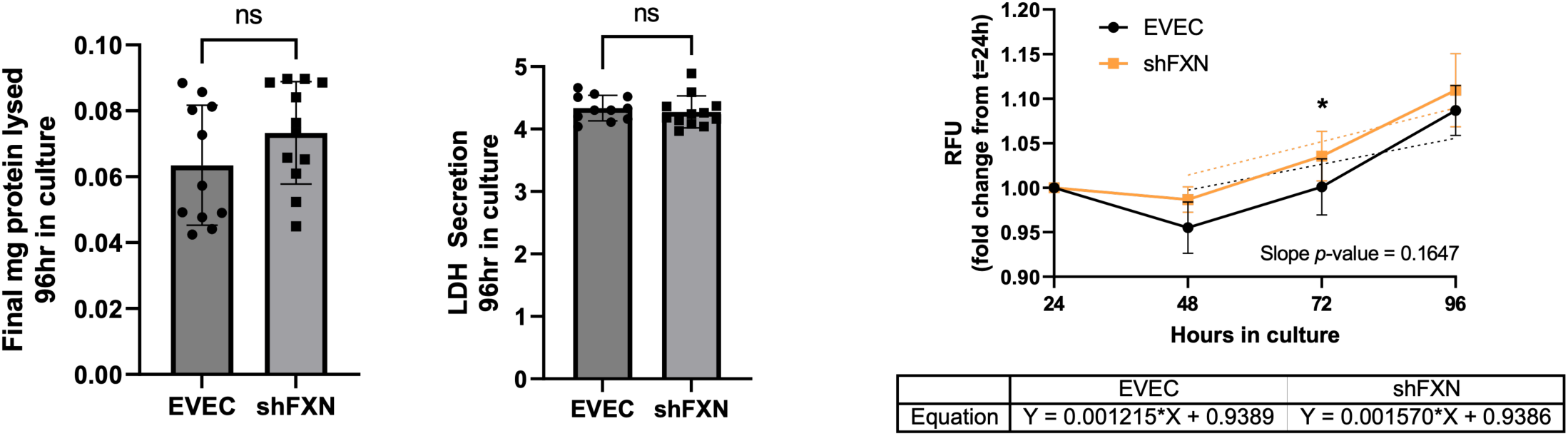
shFXN permeability values are not affected by changes in protein content, increased cell death, or changes in growth kinetics. (A) Transwells used for permeability experiments are lysed in NP-40 and quantified for total mg of protein at the final 96-hour timepoint of the assay. (B) Apical media aliquots are analyzed for LDH secretion at 96hr using the Cytox96 kit (Promega). (C) hBMVEC are plated at 5,000 cells per well and quantified for nuclear content at 24-, 48-, 72-, and 96-hours post-plating using CyQuant Direct Red (ThermoFisher). All following days are normalized to the starting concentration at 24hr for a growth curve. The linear portion of the curves (48-96hr) are analyzed via linear regression (shown with dotted lines), with statistical comparison of slope values. Student’s T-test α =0.05; ns = not significant, * p < 0.05. (A, B), EVEC; n=11, shFXN; n=12. (C) EVEC and shFXN; n= 16.

## Non-Standard Abbreviations

(CAR): Cortical Actin Ring
(EVEC): Empty Vector Endothelial Cells
(hBMVEC): Human Brain Micro-vascular Endothelial Cells
(LOFA): Late Onset Friedreich’s Ataxia
(LY): Lucifer Yellow
(OXPHOS): Oxidative Phosphorylation
(pFA): Point-Mutation Friedreich’s Ataxia
(TJ): Tight Junctions

